# The method utilized to purify the SARS-CoV-2 N protein can affect its molecular properties

**DOI:** 10.1101/2021.05.03.442392

**Authors:** Aneta Tarczewska, Marta Kolonko-Adamska, Mirosław Zarębski, Jurek Dobrucki, Andrzej Ożyhar, Beata Greb-Markiewicz

**Affiliations:** Department of Biochemistry, Molecular Biology and Biotechnology, Faculty of Chemistry, Wroclaw University of Science and Technology, Wybrzeże Wyspiańskiego 27, 50-370 Wroclaw, Poland; Department of Cell Biophysics, Faculty of Biochemistry, Biophysics and Biotechnology, Jagiellonian University, Gronostajowa 7, 30-387, Cracow, Poland

## Abstract

One of the main structural proteins of Severe acute respiratory syndrome coronavirus 2 (SARS-CoV-2) is the nucleocapsid protein (N). The basic function of this protein is to bind genomic RNA and to form a protective nucleocapsid in the mature virion. The intrinsic ability of the N protein to interact with nucleic acids makes its purification very challenging. Therefore, typically employed purification methods appear to be insufficient for removing nucleic acid contamination. In this study, we present a novel purification protocol that enables the N protein to be prepared without any bound nucleic acids. We also performed comparative structural analysis of the N protein contaminated with nucleic acids and free of contamination and showed significant differences in the structural and phase separation properties of the protein. These results indicate that nucleic-acid contamination may severely affect molecular properties of the purified N protein. In addition, the notable ability of the N protein to form condensates whose morphology and behaviour suggest more ordered forms resembling gel-like or solid structures is described.

## Introduction

Severe acute respiratory syndrome coronavirus 2 (SARS-CoV-2) has recently emerged as a highly contagious and rapidly spreading virus. Currently, scientists around the world are seeking the best possible solutions to stop the ongoing global pandemic, which began in 2019 (Zhu *et al*, 2020). SARS-CoV-2 belongs to the *Coronaviridae* family, which comprises enveloped viruses with a long, positive-sense single-stranded RNA genome. The sequenced SARS-CoV-2 genome is 29,903 bp in length (GenBank no. MN908947) and encodes nearly 30 proteins, which are responsible for host cell penetration, viral genome replication, viral gene transcription, and many other functions (Marra *et al*, 2003)(Chen *et al*, 2020b). Among all these proteins, four structural proteins can be distinguished: the spike (S) protein, the nucleocapsid (N) protein, the membrane (M) protein, and the envelope (E) protein. All of these proteins are necessary for assembling a complete viral particle and enabling the SARS-CoV-2 genomic RNA to be packed into a virion measuring approximately 100 nm (Huang *et al*, 2020).

The S glycoprotein is incorporated into the viral lipid envelope and binds to human angiotensin-converting enzyme 2 (ACE2), which serves as a virus receptor and mediates virus entry into the host cell (Huang *et al*, 2020). The most abundant viral protein, the M glycoprotein, determines the shape of the envelope (Neuman *et al*, 2011) and spans the membrane bilayer (Thomas, 2020). Importantly, due to the interactions of the M protein with all other structural proteins, it is considered the primary inducer of SARS-CoV-2 assembly (Masters, 2006). The M protein was shown to stabilize the N protein-RNA complex inside the virion (Thomas, 2020) and to support the formation of the helical ribonucleoprotein (RNP) called the nucleocapsid (Masters, 2006)(Saikatendu *et al*, 2007). The E protein is the smallest structural protein and is highly expressed during virus replication. Notably, only a small part of the E protein is incorporated into the virion envelope, and the function of this protein has not been fully elucidated to date (Venkatagopalan *et al*, 2015).

The SARS-CoV-2 N protein is a 45.5-kDa protein that presents highly conserved structural features compared to all the other nucleocapsid proteins of coronaviruses (Parker & Masters, 1990)(Masters, 1992). Importantly, the identity between sequences of SARS-CoV and SARS-CoV-2 N proteins exceeds 89%. The N protein is the only protein that is capable of binding the genomic RNA of SARS-CoV-2, which initiates the formation of a nucleocapsid that protects the viral RNA. The N protein has also been reported to enhance the efficiency of subgenomic viral RNA transcription and replication (McBride *et al*, 2014), arrest host cell cycles (Li *et al*, 2005b)(Li *et al*, 2005a), inhibit interferon production (Kopecky-Bromberg *et al*, 2007), enhance cyclooxygenase-2 (COX2) production (Yan *et al*, 2006), activate the AP1 signal transduction pathway (He *et al*, 2003), and induce apoptosis (Surjit *et al*, 2004a).

The structure of the SARS-CoV-2 N protein comprises two well-defined domains: the N-terminal domain (NTD: 48 – 175 aa) and the C-terminal domain (CTD: 247 – 364 aa) (Figure 1). While the NTD is responsible for the binding of SARS-CoV-2 genomic RNA, the CTD is critical for its homodimerization and acts as an additional RNA binding region (Cubuk *et al*, 2020b). The remaining fragments of this protein (1-47 aa, 176-246 aa, 365-419 aa) are predicted to be intrinsically disordered regions (IDRs, Figure 1). Moreover, the disordered linker between the NTD and CTD contains a conserved serine-arginine (SR)-rich sequence (183-206 aa), which is considered essential for the regulation of protein function (Peng *et al*, 2008b)(Fung & Liu, 2018)(Peng *et al*, 2008a). This region is highly phosphorylated by multiple host kinases immediately after SARS-CoV-2 infection to eliminate positive charges on the protein’s surface (Bouhaddou *et al*, 2020)(Chen *et al*, 2020a). In the case of the SARS-CoV N protein, phosphorylation enables association with specific RNA helicases and promotes the transcription of long genomic RNAs (Wu *et al*, 2014). As the infection progresses and virions assemble, N protein phosphorylation is reduced, as it is no longer required (Wu *et al*, 2014)(Wu *et al*, 2009). The structural organization of the N protein, which possesses a long and flexible linker in the middle of its sequence length, enables many conformational changes which, in turn, enable tight nucleic acid binding at multiple sites (Zeng *et al*, 2020).

**Figure 1.**
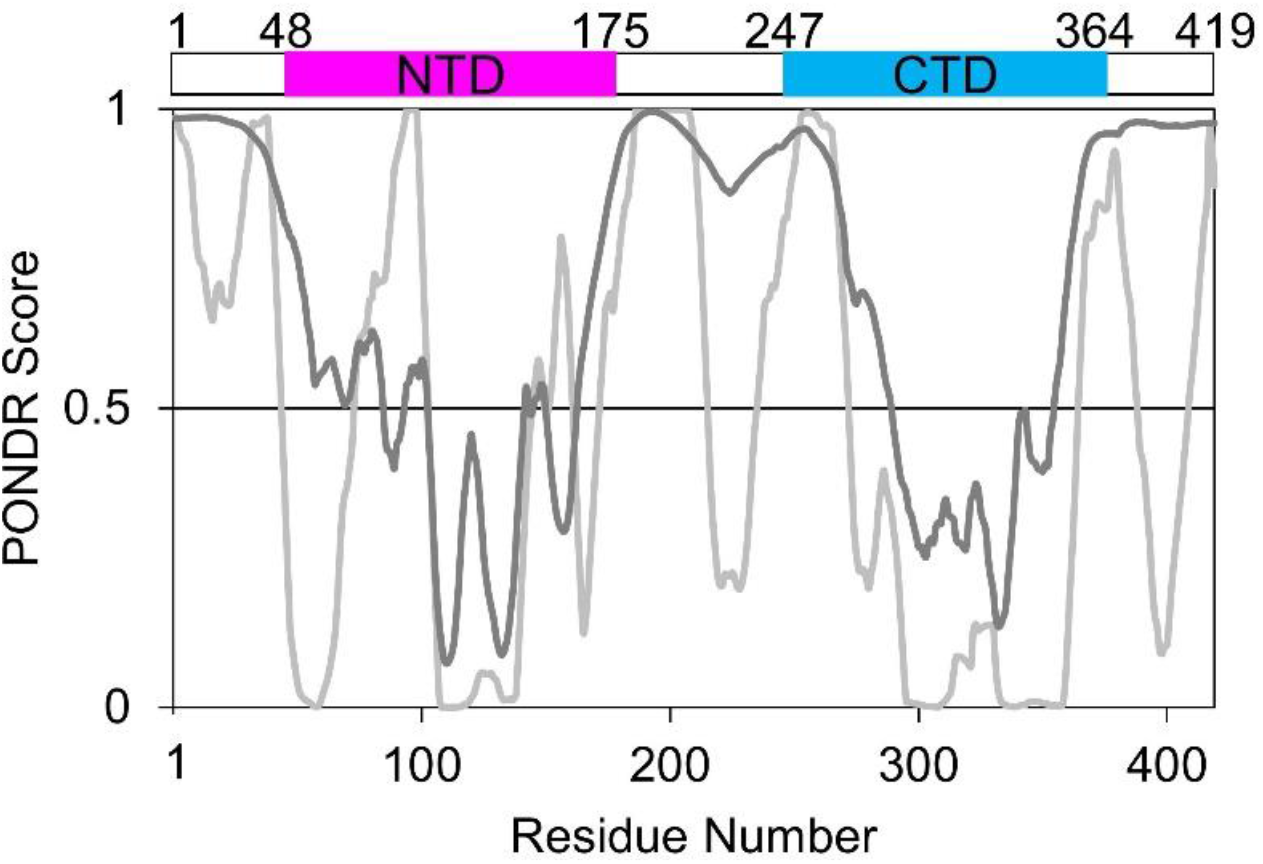
The occurrence of IDRs based on the N protein amino acid sequence. The top panel represents the localization of the NTD (magenta) and CTD (blue) domains in the N protein. The bottom panel presents PONDR-VLS2 (Peng *et al*, 2006) (dark grey line) and PONDR-VLXT (Romero *et al*, 2001) (bright grey line) disorder predictions. A score of over 0.5 indicates a high probability of disorder.

Over the last decade, scientists have focused on the consequences of spontaneously driven liquid-liquid phase separation (LLPS) in biological systems. As shown in (Lin *et al*, 2015), many RNA-binding proteins, which are characterized by the presence of long IDRs, are able to induce LLPS. This phenomenon is explained by the flexibility and multivalency of IDRs, which enables many transient interactions that promote the formation of dense, liquid condensates (Brangwynne *et al*, 2009; Holehouse & Pappu, 2018). Liquid condensates, also known as membrane-less organelles, appear to be crucial for the organization of specific cellular processes, where all necessary molecular machinery is locally concentrated to promote cellular reactions (Savastano *et al*, 2020). Such a mechanism is also considered to promote gene expression in the case of transcription (Laflamme & Mekhail, 2020). The structure and functions of the SARS-CoV-2 N protein determine its ability to induce LLPS and to interact with multiple partners (Uversky, 2013). As shown, RNA induces N protein phase separation, which probably plays a role in the specific functioning of the protein, thereby enabling proper viral replication and packaging (Iserman *et al*, 2020a; Perdikari *et al*, 2020; Savastano *et al*, 2020). Moreover, the initial site of the viral capsid and genome assembly are created in proximity to membrane structures (Stertz *et al*, 2007). Since the N protein surface is positively charged and RNA is negatively charged, condensates are driven by electrostatic effects (Iserman *et al*, 2020b).

In view of the crucial role played by the N protein, many publications that attempted to characterize this protein at the molecular level have recently been published (Ye *et al*, 2020; Zinzula *et al*, 2020; Lu *et al*, 2020; Zeng *et al*, 2020; Perdikari *et al*, 2020). These studies were conducted on recombinant N proteins isolated from bacterial cells. Considering the basic function of the protein, which is the binding of genomic RNA, we investigated whether and to what extent typical N protein isolation procedures enable a fully defined, homogeneous preparation to be obtained without contamination of nucleic acids from host cells. In this report, we present the results of a comparative structural analysis and determine to what degree the bound nucleic acids can affect the molecular properties of the N protein. We performed a reconstruction of the procedures described in recently published papers (Zeng *et al*, 2020; Perdikari *et al*, 2020; Ye *et al*, 2020; Zinzula *et al*, 2020; Lu *et al*, 2020), which resulted in protein samples that were highly contaminated with nucleic acids from the host cells used for overexpression. We believe that using contaminated samples for the study may have a significant impact on the results. Therefore, we developed a novel protocol for purifying the N protein and showed that the obtained preparation is free of nucleic acid contamination. Next, we performed comparative structural analysis of the N protein samples obtained with two protocols: a previously published protocol and a new protocol developed in our laboratory. Our aim was to determine the degree to which the presence of nucleic acids affects the molecular properties of the protein, including its propensity for oligomerization and LLPS. We demonstrate that the N protein completely separated from nucleic acids can only assemble into well-defined dimers. Our results indicate that highly oligomeric structures, described previously in the literature (e.g., tetrameric and octameric structures), could result from the presence of nucleic acids, which serve as threads to the N protein dimers. We present data indicating that the N protein contaminated with bacterial DNA/RNA drove spontaneous LLPS at a low salt concentration, while the N protein devoid of contaminating nucleic acids did not. Notably, the N-protein samples, regardless of the purification protocol used, generated specific condensates. The morphology and behaviour of these condensates suggested that they do not have a typically liquid character but instead are more structured forms (gel-like or solids). We believe that the relationship described in this report between the results of structural analyses and the method that was utilized to purify the recombinant N protein is important and should be taken into account in future analyses performed on bacterially expressed N protein.

## Results

### Elaboration of the purification procedure for obtaining nucleic acid-free protein N

The aim of this study was to obtain a homogeneous N protein preparation devoid of any nucleic acid contamination and to compare the molecular properties of that preparation with the properties of the preparation obtained using common methods. To this end, we reconstructed the procedures described in recently published papers (Zeng *et al*, 2020; Perdikari *et al*, 2020; Ye *et al*, 2020; Zinzula *et al*, 2020; Lu *et al*, 2020) (Protocol 1, Figure 2A) and developed a novel protocol (Protocol 2, Figure 4A) enabling the removal of bound nucleic acids. As the expression vector, we used pET-SUMO (EMBL, Germany), which introduces a 6xHis tag followed by a SUMO protein at the protein’s N-terminus. The 6xHis tag enables affinity chromatography purification, while the SUMO protein is expected to increase the target protein’s solubility and stability (Butt *et al*, 2005). SDS-PAGE analysis indicated the successful expression of the recombinant 6×His-SUMO-N protein (Figure 2C, T, lane 1) and its presence in the soluble fraction (Figure 2C, S, lane 2). The molecular mass (MM) of the fusion protein, determined based on the migration pattern of the Unstained Protein Weight Marker (UPMM), was determined to be 66 kDa. This measured MM was slightly higher than the theoretical MM calculated with ProtParam (59.1 kDa), which was expected, given the highly disordered character of the N protein. The purification procedure began with affinity chromatography on Ni^2+^-NTA resin. After loading of the sample, proteins unbound to the resin were washed away (Figure 2C, FT, lane 3). The 6×His-SUMO-N protein that was bound to the resin was digested with SUMO hydrolase dtUD1 (EMBL, Germany)(Weeks *et al*, 2007) on the column. As the 6×His tag was removed from the digested N protein, it was collected as a flow-through (Figure 2C, FT_N,_ lane 4). The remaining proteins were eluted from the resin with buffer containing 500 mM imidazole. The digestion step was determined to be efficient, as the N protein was not present in that fraction (Figure 2C, elu, lane 5).

**Figure 2.**
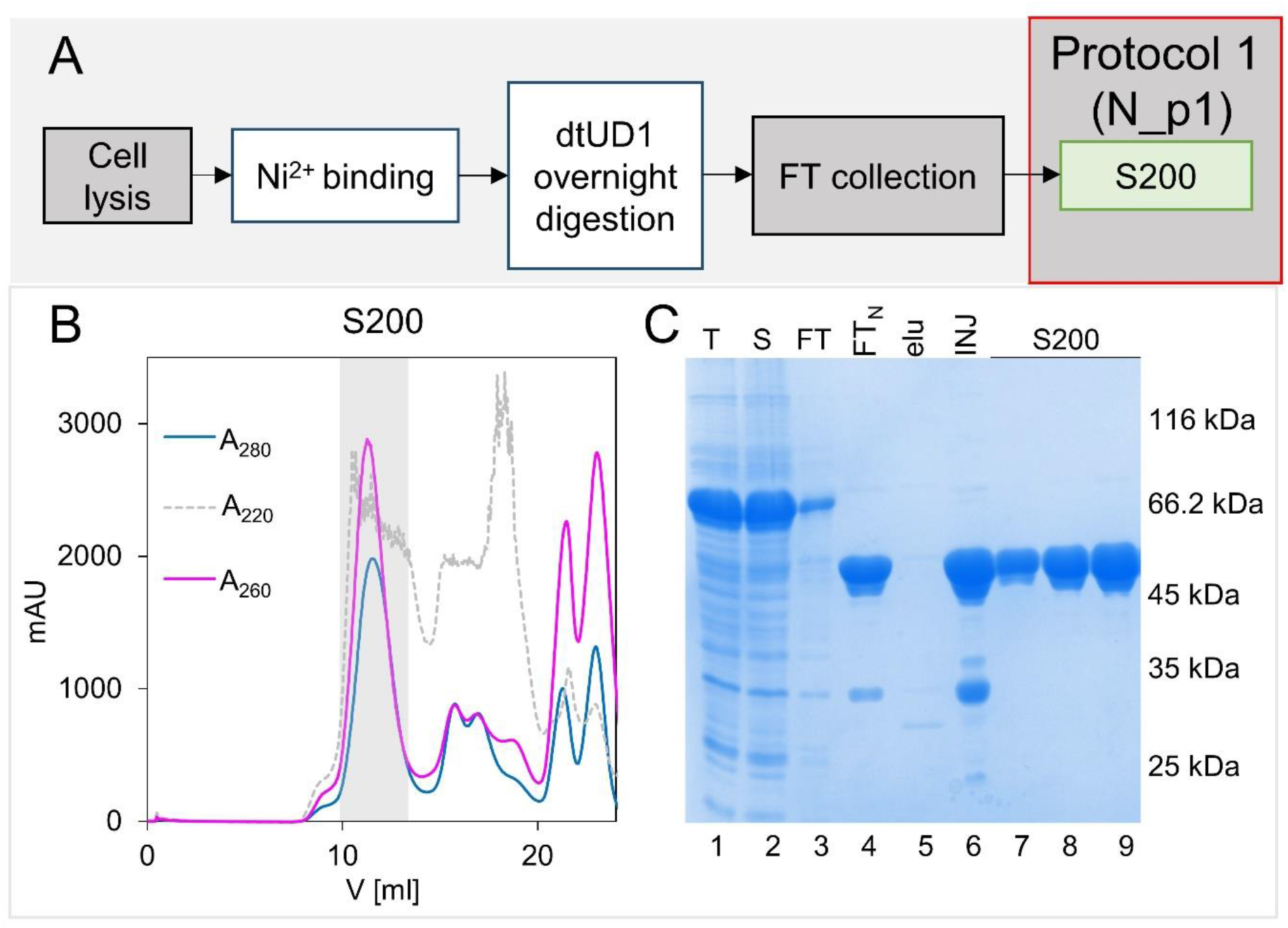
N protein purification - Protocol 1. A) schematic representation of the N protein purification process according to Protocol 1. For more details, see the Materials and Methods section. B) Preparative SEC on the Superdex 200 column. Fractions containing the N protein are marked with the grey colour. C) BlueStain Sensitive-stained SDS-PAGE analysis of the N protein samples presenting the purification procedure. Lane 1 - T, the total bacterial protein fraction; lane 2 - S, the soluble proteins fraction; lane 3 - FT, the fraction containing proteins not bound to the Ni2+-NTA resin; lane 4 FTN, proteins washed from the resin after digestion with dtUD1protease; lane 5 - elu, fractions eluted from the Ni2+-NTA resin; lane 6 - INJ, combined fractions injected on the Superdex 200 column; lanes 7-9, the main peak from the Superdex 200 column, containing N protein.

### Protocol 1

Collected fractions containing the N protein were concentrated and loaded on a Superdex 200 column (Figure 2B, C, INJ, lane 6). The purity determination of the obtained N protein sample (N_p1), which based on SDS-PAGE analysis (Figure 2C, lanes 7-9), revealed that the sample was over 95% pure. However, during spectroscopic determination of the protein concentration, we observed a shifting of the absorbance maximum, which is usually expected at approximately 280 nm, towards shorter wavelengths. For this reason, we decided to determine if the purified protein sample had been contaminated by the nucleic acids of the host cells (*E. coli*). To this end, the N_p1 sample was incubated at 50°C in the absence or presence of SDS. All the samples were analysed on agarose gels containing a specific nucleic acid dye. As presented in Figure 3A, both the control sample (lane 2) and the sample after incubation at 50°C (lane 3) presented a fluorescence signal in the loading wells, which indicated the presence of nucleic acids. The fact that the nucleic acids did not migrate in the electric field suggests that they either form large molecular complexes that are not able to enter the gel pores or that the nucleic acid’s negative charges are partially neutralized by binding to the protein. In the case of the N_p1 sample incubated at 50°C in the presence of SDS, the result was different (Figure 3A, lane 4). This strong anionic detergent forms complexes with the protein’s backbone, thereby adding negative charges to the protein’s molecule, which interrupts the electrostatic interactions between the protein and nucleic acid. Free nucleic acids entered the gel and migrated in the electric field. Notably, two subpopulations of nucleic acids on the gel were observed (Figure 3A, lane 4). The first population migrated as a defined band, while the other migrated as a smear.

**Figure 3.**
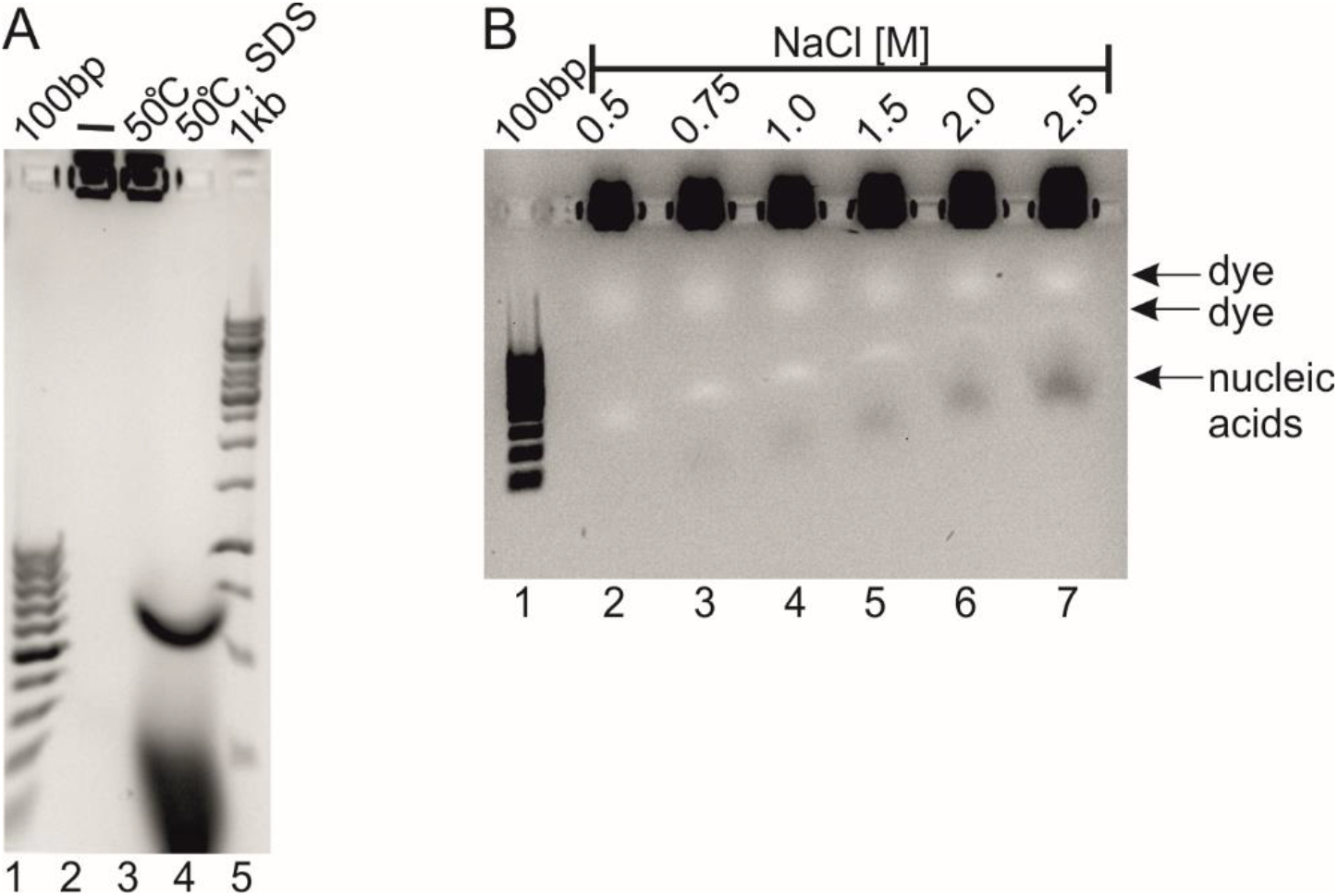
Agarose gel electrophoresis of the N protein samples obtained with Protocol 1. A) Detection of nucleic acids in the N_p1 sample. Lane 1 (100bp), DNA ladder; lane 2 (-), the untreated N_p1 sample; lane 3 (50°C), the N_p1 sample incubated at 50°C; lane 4 (50°C, SDS), the N_p1 sample incubated with SDS at 50°C; lane 5, (1kb), DNA ladder. B) The N_p1 sample in the presence of an increasing concentration of NaCl. Lane 1 (100bp), DNA ladder, lane 2-7, the N_p1 sample incubated in a buffer supplemented with various concentrations of NaCl. The concentration of NaCl in the protein samples is provided at the top of the image.

This result motivated us to optimize the purification method and to develop a protocol enabling N proteins not contaminated with nucleic acids to be obtained. First, we decided to observe the impact of increasing ionic strength on the stability of the protein-DNA complex. To this end, the N_p1 protein was incubated in a buffer containing different concentrations of sodium chloride. As presented in Figure 3B, sodium chloride at a concentration below 1.5 M had no impact on the complexes, whereas concentrations of sodium chloride of 1.5 M and higher had minor impacts on the stability of the complexes. Only a small fraction of the nucleic acid was observed in a free form, while most of the complex remained intact. These results indicated that the N protein can form very tight complexes with bacterial, nonspecific nucleic acids. Based on this observation, we decided to significantly modify the purification method and to introduce an additional step: chromatography on a Sepharose resin with immobilized heparin. Heparin is highly sulfonated glycosaminoglycan. The anionic surface of heparin can mimic negatively charged nucleic acids. We previously employed this approach for the efficient purification of the DNA binding domains of nuclear receptor transcription factors (Niedziela-Majka *et al*, 2000).

### Protocol 2

As described above, 6×His-SUMO-N was loaded onto Ni^2+^-NTA resin, digested on the column, and collected as flow-through (FT) (Figure 4A). Next, the collected fractions were loaded onto a system of two connected in-series 1 ml Heparin Sepharose columns (Figure 4C, INJ, lane 1). Although the amount of injected protein was relatively low, only this extension of the column bed resulted in the expected and efficient elimination of nucleic acids, which did not bind to the column and were collected as FT (Figure 4B, fractions A5, A12). SDS-PAGE and agarose gel electrophoretic analysis revealed that the fraction unbound to the Heparin Sepharose columns (fractions A5 and A12) did not contain the target protein (Figure 4C, lanes 3 and 4). The applied linear gradient of sodium chloride up to 1 M enabled the elution of the proteins that were bound to the resin (Figure 4B, C lanes 5, 6). SDS-PAGE analysis showed that the N protein was eluted from the columns with buffer supplemented with 0.7 M sodium chloride (Figure 4C, lane 6). The apparent MM (50 kDa) of the protein, as determined based on the relative migration to the migration pattern of the UPMM (Figure 4C, lane 2), differs from the MM of the theoretical N protein (45.6 kDa). The difference in MM can be explained by the presence of disordered regions (Receveur-Bréchot *et al*, 2006) in the structure of the N protein. Although the purity of the obtained protein sample that was examined by SDS-PAGE electrophoresis was high, agarose gel electrophoresis revealed the presence of nucleic acid impurities in eluted fractions (see Figure 5B, lane 2). Similar to the N_p1 sample, the nucleic acids remained in the loading wells, indicating the presence of large complexes of the N protein with nucleic acids. Importantly, the nucleic acid signal was considerably lower in comparison to the N_p1 sample. To remove these residual impurities, we decided to take the additional purification step of utilizing tandem SEC chromatography on two connected Superdex 200 columns. The eluted fractions were collected in 0.25-ml volumes, and spectroscopic and electrophoretic analyses were subsequently performed. In the elution profile, which was monitored by the absorbance at 280 and 260 nm, two peaks could be distinguished (Figure 4D). Notably, a shift of the absorbance maximum position for the eluted fractions was observed. For the initial fractions eluted from the SEC columns (Figure 5A, A10-B1), the absorbance maximum was observed at 260 nm, indicating the presence of nucleic acids. The absorption profiles of the B2 and B3 fractions (Figure 5A) were broad, and their maximum shifted towards longer wavelengths. The B4-B10 fractions (Figure 5A) collected from the second peak that was eluted from the Superdex 200 columns (see Figure 4D) presented a defined absorbance maximum at 280 nm. The electrophoretic analysis, which was performed at the same time, revealed decreasing amounts of nucleic acids in the subsequent fractions. The A10-B1 fractions contained the highest amounts of nucleic acids (Figure 5B, lanes 3-10), the B2 and B3 fractions presented moderate amounts of nucleic acids (Figure 5B, lanes 11-12), and the B4-B7 fractions were free of nucleic acids (Figure 5B, lanes 13-15). To summarize, simultaneous SEC on the two connected Superdex 200 columns enabled efficient separation of nucleic acid contamination from the purified N-protein. The fractions from the first peak eluted from the Superdex 200 columns were enriched in nucleic acids, while the fractions from the second peak contained the purified N protein and were free of such contamination. In later sections of the manuscript, this fraction is referred to as N_p2.

**Figure 4.**
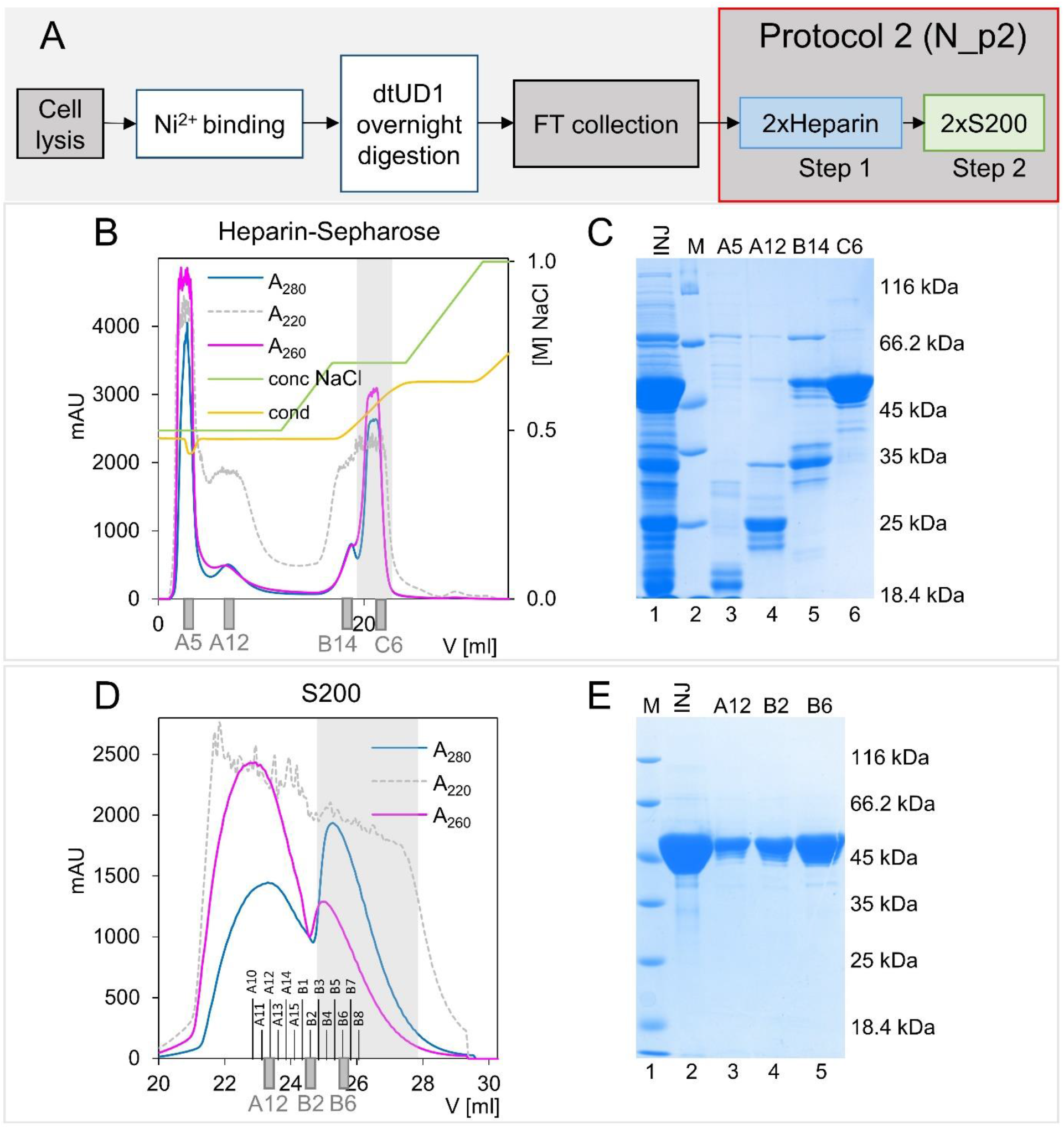
N protein purification - Protocol 2. A) Scheme of the N protein purification process according to Protocol 2. For more details, see the Materials and Methods section. B) Chromatography on the Heparin Sepharose columns. Fractions containing the N protein are marked with the grey colour. Fractions analysed with SDS-PAGE are marked under the graph. C) BlueStain Sensitive-stained SDS-PAGE analysis of the N protein samples presenting the results of Heparin Sepharose column purification step. Lane 1 - INJ, combined fractions loaded on Heparin Sepharose column; lane 2 - M, molecular weight marker; lanes 3-6, elution profile. D) Preparative SEC on two connected in series Superdex 200 columns. Fractions containing the N protein are marked with the grey colour. Fractions analysed with SDS-PAGE are marked under the graph. E) BlueStain Sensitive-stained SDS-PAGE analysis of the N protein samples presenting the SEC purification step. Lane 1 - M, molecular weight marker; lane 2 - INJ, combined fractions eluted from the Heparin-Sepharose column, loaded on the Superdex 200 columns; lanes 3-5, elution profile.

**Figure 5.**
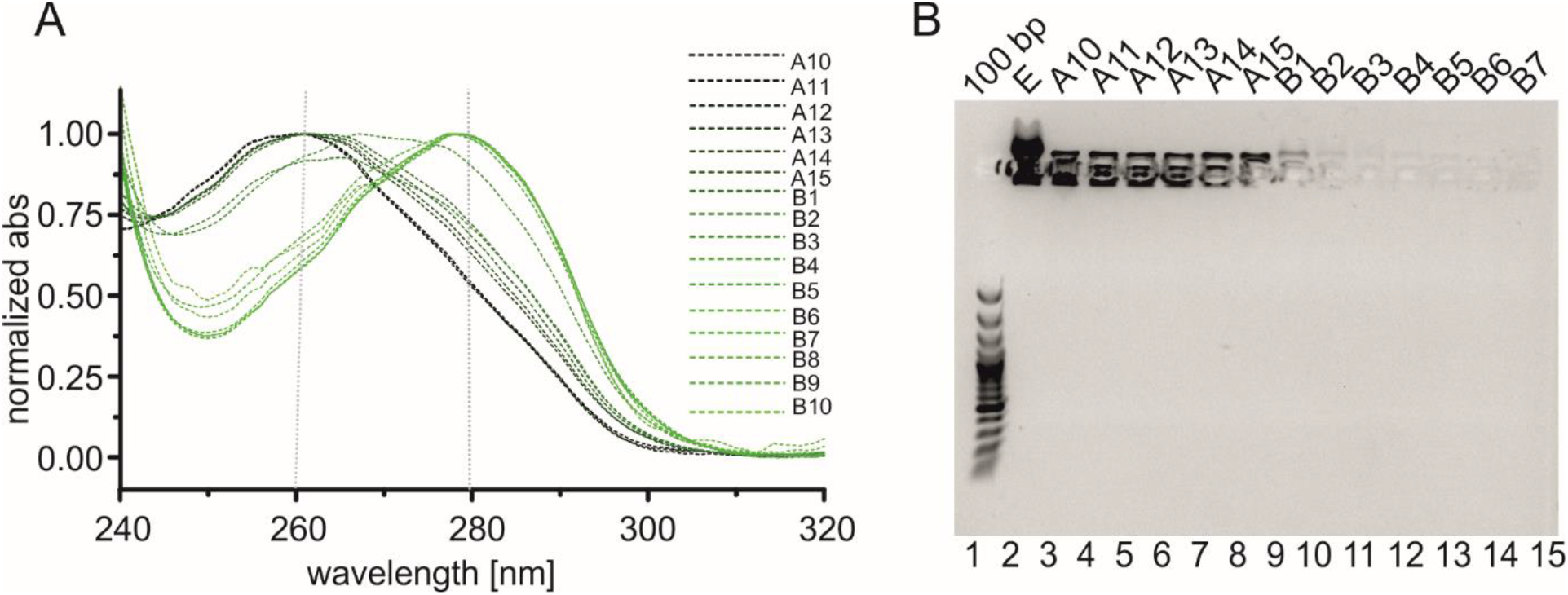
Spectroscopic and electrophoretic analysis of the fractions eluted from the tandem SEC. A) Normalised absorption profiles of the eluted samples (A10-B10) obtained with NanoDrop; B) The agarose gel stained with SimplySafe dye. Lane 1, (100 bp) DNA ladder; lane 2-15, fractions A10-B7 eluted from the Superdex 200 columns.

### Impact of the presence of nucleic acids on the secondary structure of the N protein

Available data suggested that the N protein forms partially disordered dimers and higher oligomers (Zeng *et al*, 2020)(Perdikari *et al*, 2020). Based on our observations, we hypothesized that previously published biochemical analyses of the N protein might contain some errors resulting from the overlooked presence of nucleic acids derived from recombinant expression host cells. To test this hypothesis, we first performed far-UV circular dichroism (CD) spectroscopy analysis of the N_p1 and pure N_p2 samples. CD spectroscopy is a commonly utilized method for determining a protein’s secondary structure content and folding properties (Kelly & Price, 1997). Moreover, nucleic acids are able to absorb circularly polarized light (Gray *et al*, 1995). The spectra obtained for both the contaminated (N_p1) and free (N_p2) nucleic acid samples showed similar optical properties in the range of 210-240 nm (Figure 6), indicating that the secondary structure content of the N protein does not change in the presence of nucleic acids. The spectral deconvolution revealed that over 50% of the protein exists in a disordered conformation (Figure 6, inset). Notably, the N protein gives off a signal with a maximum at 236 nm that absorbs left circularly polarized light, indicating the presence of a unique optically active chiral centre. The active chiral centre is preserved after the protein forms complexes with nucleic acids. However, the N_p1 and N_p2 samples presented different optical properties between 250 and 300 nm. In this wavelength range, the N_p1 sample showed a positive signal with a maximum at 264 nm (Figure 6), which is characteristic of the presence of nucleic acids (Miyahara *et al*, 2016)(Kypr *et al*, 2012). In addition, this spectral shape is typical for RNA (Miyahara *et al*, 2016)(Kypr *et al*, 2012).

**Figure 6.**
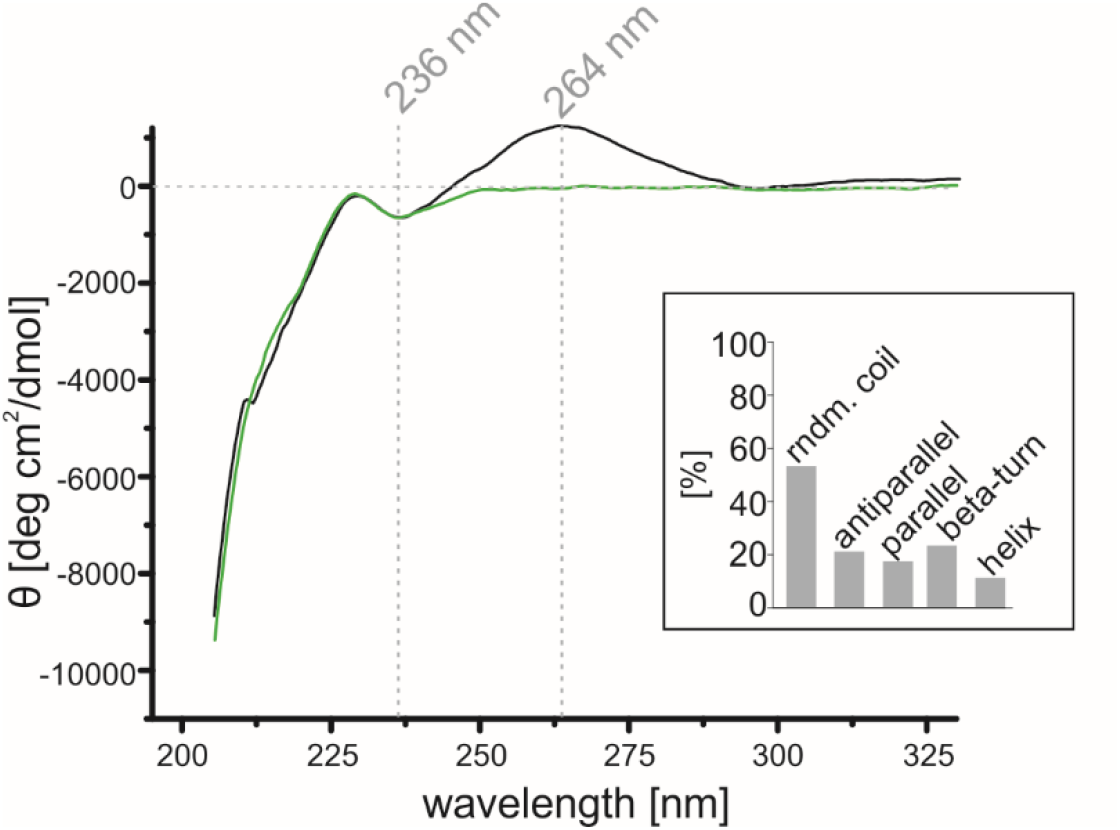
CD spectra of the N protein. The UV spectra obtained for the N_p1 (black line) and N_p2 (green line) samples. Both spectra display a strong negative signal with the maximum close to 200 nm, which corresponds to the presence of a disordered conformation. Specific negative maxima are also present in both samples at 236 nm. The N_p1 sample shows a positive signal with the maximum at 264 nm, which corresponds to the presence of nucleic acids. The inset presents the estimated percentage content of the secondary structures obtained from the deconvolution of the averaged N_p2 spectrum at a range of 210-230 nm. The deconvolution was conducted using CDNN 2.1 software (available at: (http://www.gerald-boehm.de/).

### Identification of the type of nucleic acids

It is known that *Coronaviridae* are single-stranded RNA viruses (Payne, 2017) and that their nucleocapsid proteins have an intrinsic ability to bind nucleic acids (McBride *et al*, 2014). The composition of nucleic acids that form complexes with the recombinant N protein expressed in *E. coli* cells is clearly different from that which interacts with the N proteins during infection. Nevertheless, according to our observations, the strength of the interaction between the expressed N protein and bacterial nucleic acids is very strong. Notably, agarose gel electrophoresis of N_p1 revealed the presence of two subpopulations of nucleic acids in the samples. For this reason, we decided to determine more precisely the types of nucleic acid interacting with the protein. To this end, the N protein samples purified according to Protocol 1 (N_p1), which were contaminated with bacterial nucleic acids, were first treated with either RNase A or DNase I and analysed electrophoretically. Although the cleavage reaction was performed under optimal conditions, a strong nucleic acid-specific signal was still observed in the loading wells (Figure 7A, lanes 2,3). This result indicates that the N_p1-complexed nucleic acids are protected from the activities of nucleases. Therefore, we decided to perform a similar experiment with the fraction deprived of the N protein during purification on Heparin Sepharose columns. This sample was referred to as sample (S) (see Figure 4B, C, fraction A5) and contained a high amount of nucleic acids. As the control sample (CS), fraction C6 containing the N protein eluted from the Heparin Sepharose resin (see Figure 4B, C) was used.

**Figure 7.**
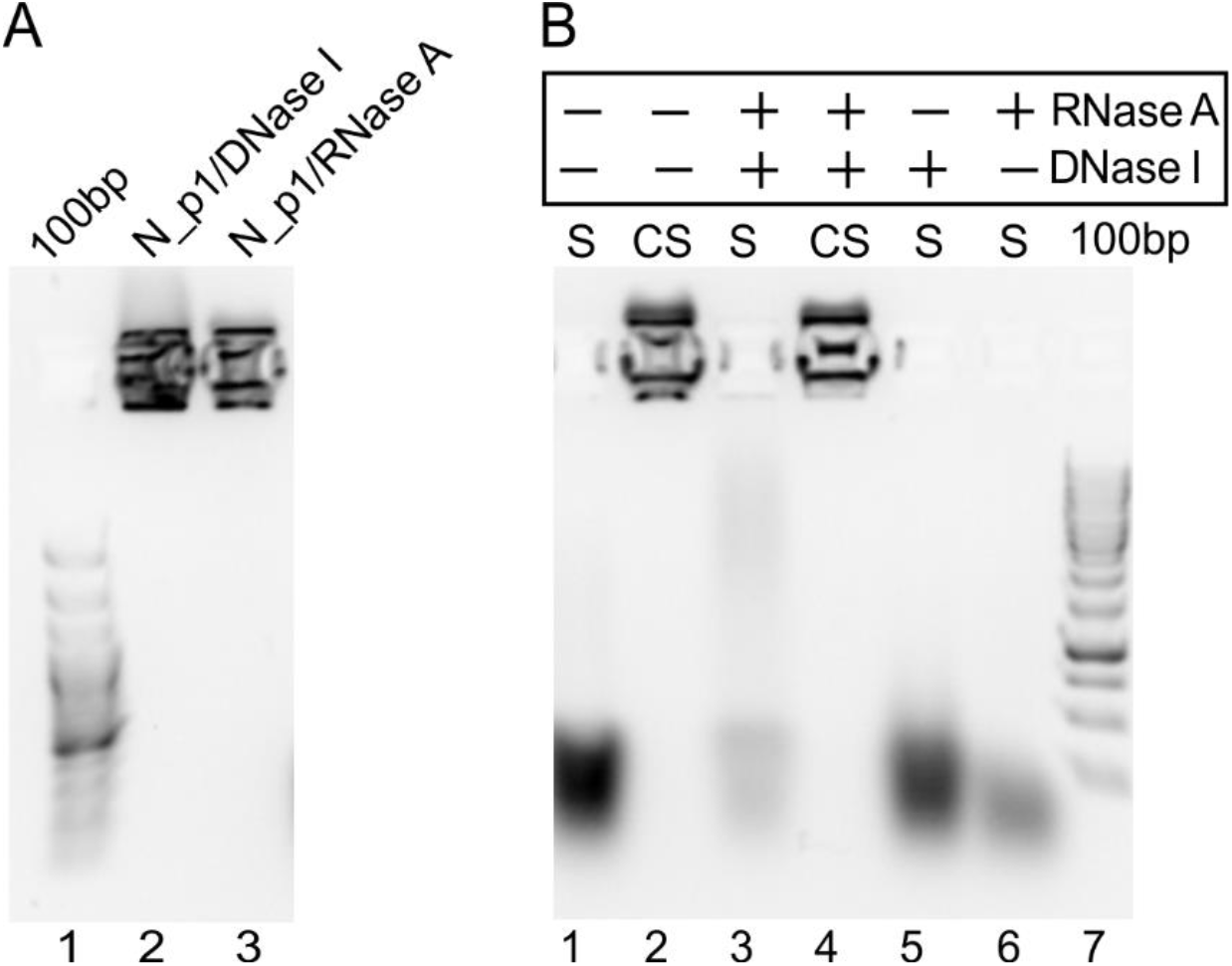
Determination of the type of nucleic acids bound to the N protein. A) Electrophoretic analysis of the N protein sample purified according to Protocol 1. Lane 1, DNA ladder; lane 2, N_p1 protein digested with DNase I; lane 3, protein digested with RNase A. B) Electrophoretic analysis of the fractions obtained during Heparin Sepharose chromatography. Lane 1, the unbound to the resin sample (S) containing freed nucleic acids; lane 2, the control sample (CS) containing the C6 fraction eluted from the Heparin Sepharose column; lanes 3 and 4, S and CS digested with both DNase I and RNase A; lane 5, S digested with DNase I; lane 6, CS digested with RNase A; lane 7, DNA ladder.

First, both samples (S and CS) were cleaved with DNase I (Figure 7B, lane 3) and RNase A (Figure 7B, lanes 4). While the nucleic acids present in the S were completely hydrolysed, in the CS sample, the nucleic acids forming a complex with the N protein were protected from nucleases (Figure 7A). The analysis of the digestion products of the nucleic acids present in the S sample yielded interesting results. Digestion with DNase I resulted in a band with moderate intensity (Figure 7B, lane 5), which indicates that most of the nucleic acid population was not a substrate for this nuclease. In the case of digestion with RNase A, the band intensity was significantly decreased (Figure 7B, lane 6). However, the nucleic acids were not completely hydrolysed. Taken together, these results indicate that the N protein binds bacterial RNA but that it also has the ability to bind DNA.

### Impact of the presence of nucleic acids on the quaternary structure of the N protein

As shown previously, the N protein possesses a quaternary structure (Zeng *et al*, 2020). It was reported that the N protein can form dimeric structures, but the presence of higher oligomers was also suggested (Zeng *et al*, 2020)(Perdikari *et al*, 2020)(Ye *et al*, 2020). To verify the impact of the presence of nucleic acids on the quaternary structure of the recombinant N protein, we analysed its hydrodynamic properties. First, we performed analytical SEC to compare the elution profiles of the N_p1 and N_p2 samples. In the case of the sample containing nucleic acids (N_p1), two peaks were observed: the first (A12 fraction) at an elution volume of 11.8 ml (corresponding to 104 kDa) and the second (B8 fraction) at a volume of 14.5 ml (corresponding to 23 kDa) (Figure 8A). Importantly, the absorbance signal of the first peak was higher for the 280-nm wavelength. In contrast, for the second peak, higher absorbance was observed at 260 nm (Figure 8A). SDS-PAGE analysis of the injected sample (Figure 8B, INJ, lane 1) and selected fractions revealed that the N protein was only present in the first peak (Figure 8B, fraction A12, lane 3), with it being absent in the B8 fraction (Figure 8B, fraction B8, lane 4) from the second peak. During the analytical SEC, some of the remaining nucleic acids that were bound to the protein dissociated from the protein. In the case of the N protein sample purified according to protocol 2 (N_p2), only one peak, corresponding to 104 kDa, was observed (Figure 8A). The N protein free from nucleic acid contamination was eluted at the elution volume corresponding to a dimer.

**Figure 8.**
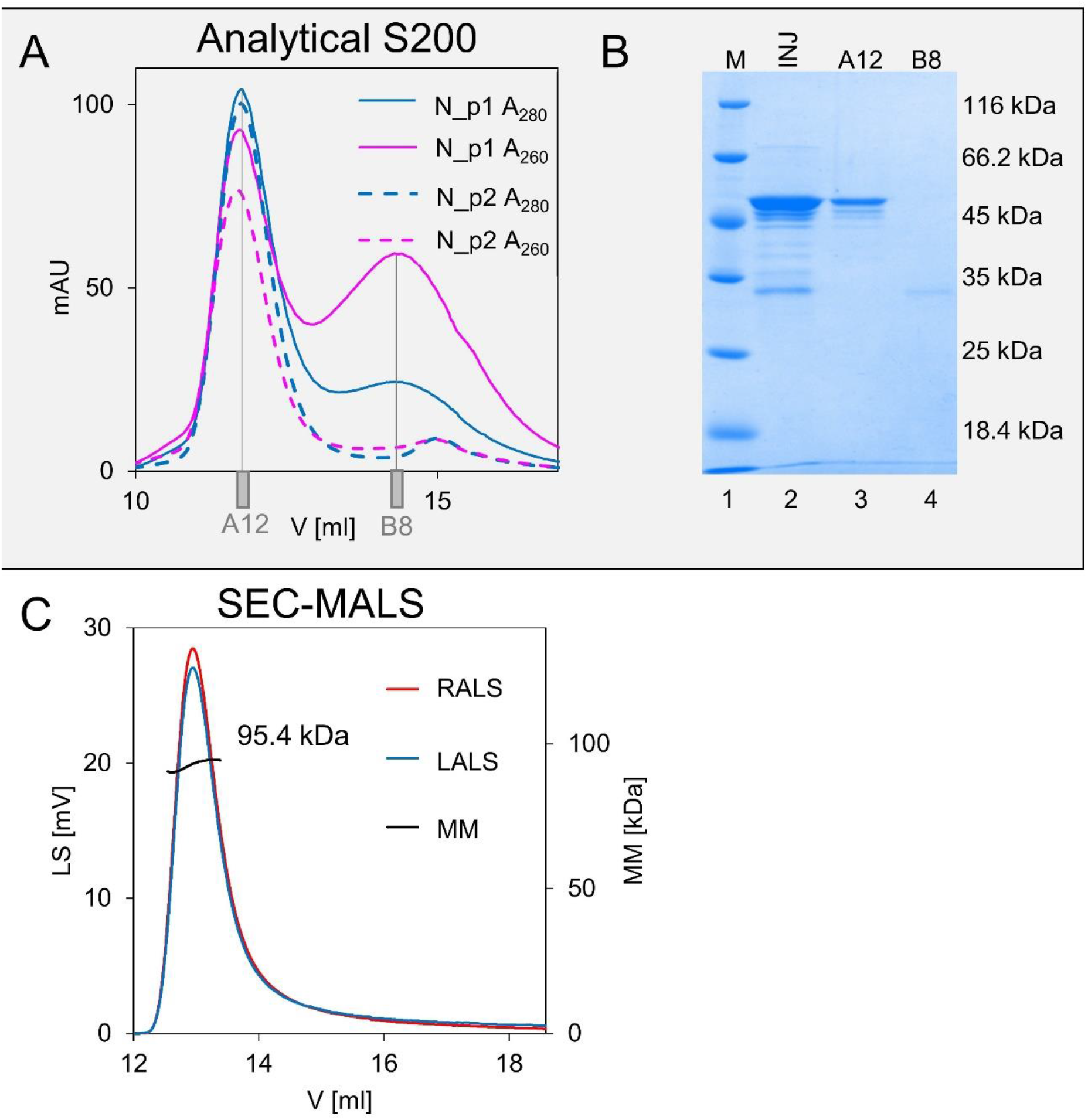
Hydrodynamic properties of the purified N protein. A) Analytical SEC profile for the N_p1 (solid lines) and N_p2 (dashed lines) samples. The blue colour represents absorbance at 280 nm, and the pink colour represents absorbance at 260 nm. B) SDS-PAGE analysis of the N_p1 samples. Lane 1, molecular weight marker; lane 2 - INJ, N_p1 fraction injected on the column; lane 3, first peak (A12, 11.8 ml); lane 4, second peak (B8,14.5 ml). C) SEC-MALS profile of the N_p2 sample.

The results of the analytical SEC were additionally confirmed by the SEC-MALS experiment (Figure 8C). The elution profile and MM (95.4 kDa) determined from right angle light scattering (RALS) and left angle light scattering (LALS) signals are in keeping with the SEC results and confirm the existence of the N protein in a dimeric form in the N_p2 sample. In the case of the N_p1 sample, the corresponding MM of the protein could not be determined. These difficulties resulted from the overestimated concentration value of the protein sample. The measured value of the absorbance at 280 nm was affected by the presence of nucleic acids, with the maximum absorbance being observed at 260 nm. Nevertheless, the experiment revealed, in divergence from the results obtained with the monodispersed N_p2 population, that N_p1 forms more heterogeneous populations that correspond to higher oligomers (Figure EV1).

To determine the ability of the N protein to form higher oligomers, we applied sedimentation velocity analytical ultracentrifugation (SV-AUC). The experiment was performed for the protein at three different concentrations and revealed that the fully purified N_p2 sample sediments as a homogeneous population (Figure 9A). The water-standardized sedimentation coefficient (s_20,w_) values were 3.71, 3.78 and 3.79 s for the analysed protein concentrations: 2.5, 0.5 and 0.75 mg/ml, respectively (Table 1). The mean calculated MM for the three analysed concentrations was 84.07 ± 1.62 kDa, which corresponds to the theoretical MM value of the dimer (91.1 kDa). In the case of the protein contaminated with nucleic acids (N_p1), interpretable values were only obtained for the analysed protein concentrations: 0.25 and 0.5 mg/ml (Figure 9B). The analysis of the protein samples at two concentrations indicated the existence of two subpopulations (Figure 9B). The mean calculated s_20,w_ value of the first population (3.695 ± 1.34) corresponds to the mean s_20,w_ value obtained for the N_p2 sample (3.76 ± 0.04). The second population indicates the presence of larger molecules. Their s_20,w_ values were 5.3 and 5.1 s for N_p1 analysed at 0.25 and 0.5 mg/ml concentrations, respectively (Table 1). Because it is not possible to determine the exact concentration of a protein in a complex with nucleic acids, the calculation of the molecular weight values cannot be precise. However, the SV-AUC experiment revealed that the N protein occurs as a dimer and that larger species mimicking higher oligomers can only be formed in the presence of nucleic acids.

**Table 1.** Hydrodynamic properties of the N protein samples determined by analytical ultracentrifugation.

|  | peak I |  |  |  |  | peak II |  |  |
| --- | --- | --- | --- | --- | --- | --- | --- | --- |
| | C<br>[mg/ml] | $S_w$ [S]<br>[S] | $S_{w(20,w)}$<br>[S] | MM<br>[kDa] | $R_s$<br>[nm] | $S_w$ [S]<br>[S] | $S_{w(20,w)}$<br>[S] | MM<br>[kDa] |
| N_p1 | 0.25 | 3.65 | 3.79 | 78.2 | 5.12 | 5.3 | 5.5 | 135 |
|  | 0.5 | 3.5 | 3.6 | 50 | 3.42 | 5.1 | 5.3 | 89 |
|  | 0.75 | 3.6 | 3.76 | 55 | 3.6 | 5.2 | 5.4 | 95 |
| N_p2 | 0.25 | 3.43 | 3.71 | 85 | 5.32 |  |  |  |
|  | 0.5 | 3.50 | 3.78 | 85.2 | 5.38 |  |  |  |
|  | 0.75 | 3.50 | 3.79 | 85 | 5.19 |  |  |  |

**Figure 9.**
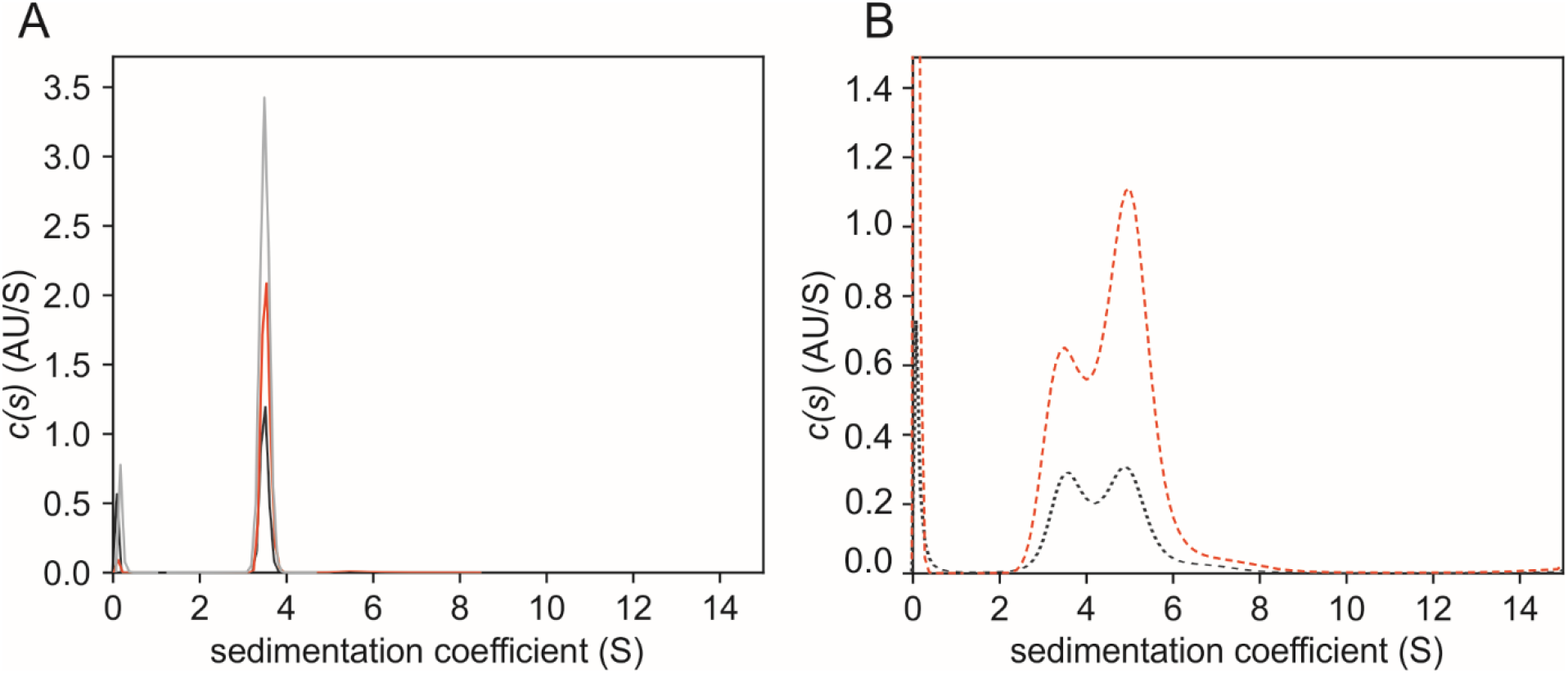
SV-AUC analysis of the N protein. Each graph presents a coefficient distribution (c(s)) model of SEDFIT from the SV data obtained for different concentrations of samples: 0.25 (black line), 0.5 (red line) and 0.75 mg/ml (grey line) of samples. A) N_p2 samples; B) N_p1 samples.

### Propensity of the N protein for LLPS

The N protein is also the RNA binding protein. Moreover, the ability of the N protein to induce spontaneous LLPS has already been shown (Cubuk *et al*, 2020a)(Lu *et al*, 2020). However, based on the results that we obtained through oligomerization studies, we investigated whether the purification procedure could also affect the LLPS propensity of the recombinant N protein. To this end, we decided to perform comparative analysis of the ability to induce LLPS of the recombinant N protein purified according to protocols 1 and 2. To set the uniform methodology for the comparative studies, we microscopically examined different concentrations of the N_p1 and N_p2 samples in HEPES buffer supplemented with 500 mM NaCl. We found that the N_p1 and N_p2 samples at concentrations up to 10 mg/ml remained in a one-phase solution (Figure 10A), and we therefore used this concentration as the initial condition for preparing the defined samples. It was previously shown that the N protein promotes LLPS in low ionic strength (Perdikari *et al*, 2020)(Cubuk *et al*, 2020a). Therefore, we tested the effect of ionic strength on the ability of the N protein to provoke LLPS. The nucleic acid-contaminated protein (N_p1) was analysed at a concentration of 1 mg/ml, whereas N_p2 was analysed at concentrations of 1 and 2 mg/ml. As presented in Figure 10A and 10B, the N_p1 protein sample (contaminated with bacterial nucleic acids) was able to form condensates in buffers of 250 mM NaCl and lower. The formed condensates were round and diffused in the solution. Notably, several minutes after formation, the condensates settled on the glass slide, which indicated that they were liquid condensates. The highly purified N protein (N_p2) samples with no contamination of nucleic acids were not able to provoke phase separation under the corresponding conditions (Figure 10B). This test was repeated with a higher N_p2 protein sample concentration (2 mg/ml) (Figure 10B) and in the presence of a molecular crowding mimicking agent (PEG300 and PEG8000). However, we were also not able to observe the formation of any condensates under these conditions. Our results indicated that the contamination of a sample by bacterial nucleic acids can affect the propensity of the N protein to drive LLPS.

**Figure 10.**
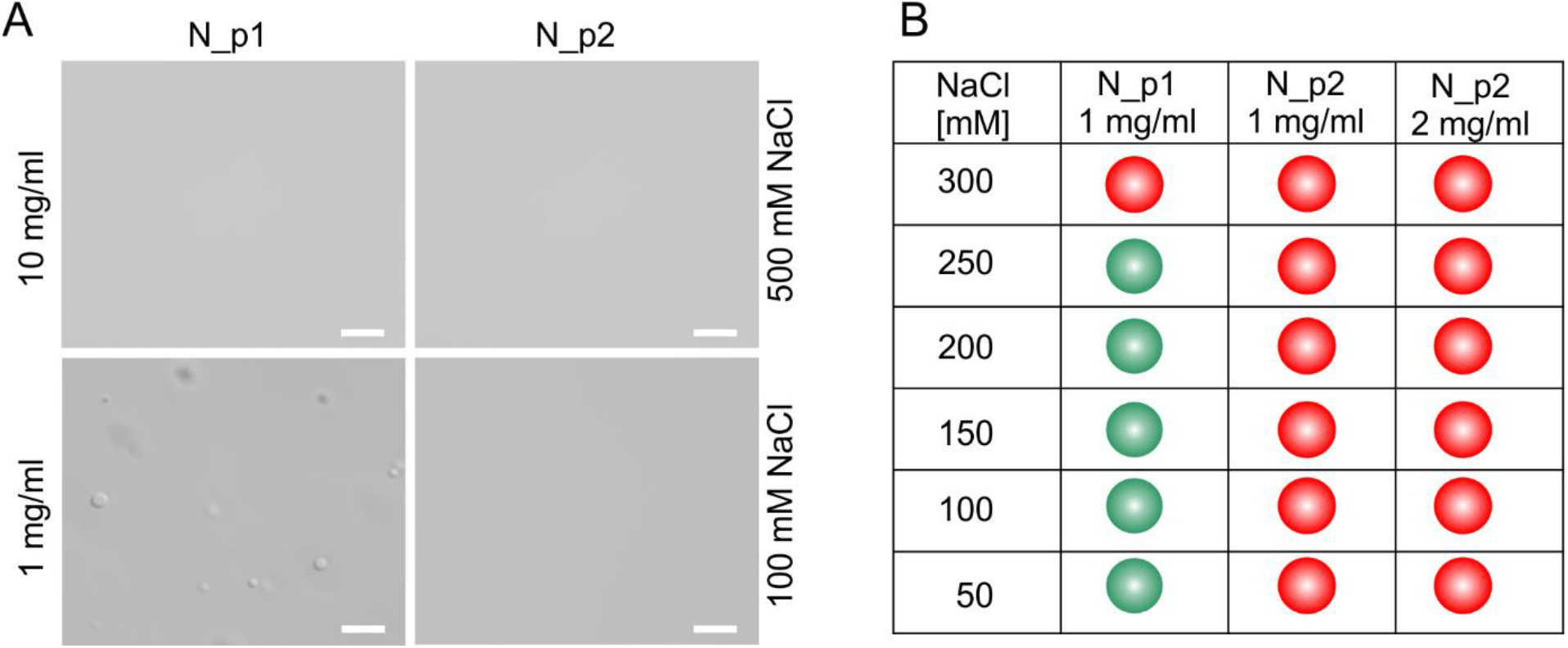
The propensity of the N protein for inducing LLPS. A) The representative images obtained by differential interference contrast (DIC) microscopy showing the concentrated (10 mg/ml) and diluted (1 mg/ml) solutions of the N protein in the buffer supplemented with high (500 mM) and low (100 mM) concentrations of NaCl. Scale bar 5 µm. B) A diagram indicating the condition at which LLPS is induced (green dots) or is not induced (red dots) by the N protein.

### N protein and other types of phase transition

Our results revealed that the recombinant N protein interacts with bacterial RNA. Therefore, we decided to analyse the propensity of the N proteins purified according to Protocols 1 and 2 to form condensates in the presence of total bacterial RNA. To this end, we used a constant concentration (1 mg/ml) of the N protein and increasing amounts of bacterial RNA (0, 25, 50, 100 and 200 ng). The N_p1 samples were only analysed in a buffer containing 300 mM NaCl, while the N_p2 samples were studied in a buffer containing both 300 mM NaCl and 150 mM NaCl. Under these conditions, condensates were not observed (see Figure 10B). The addition of RNA to the tested samples affected the protein’s properties: in all the analysed samples, regardless of the concentration of RNA, some irregular particles were observed (Figure 11). An increase in the number of particles correlated with an increase in the added RNA concentration. Notably, these forms did not resemble liquid condensates but rather appeared to be solid aggregates.

**Figure 11.**
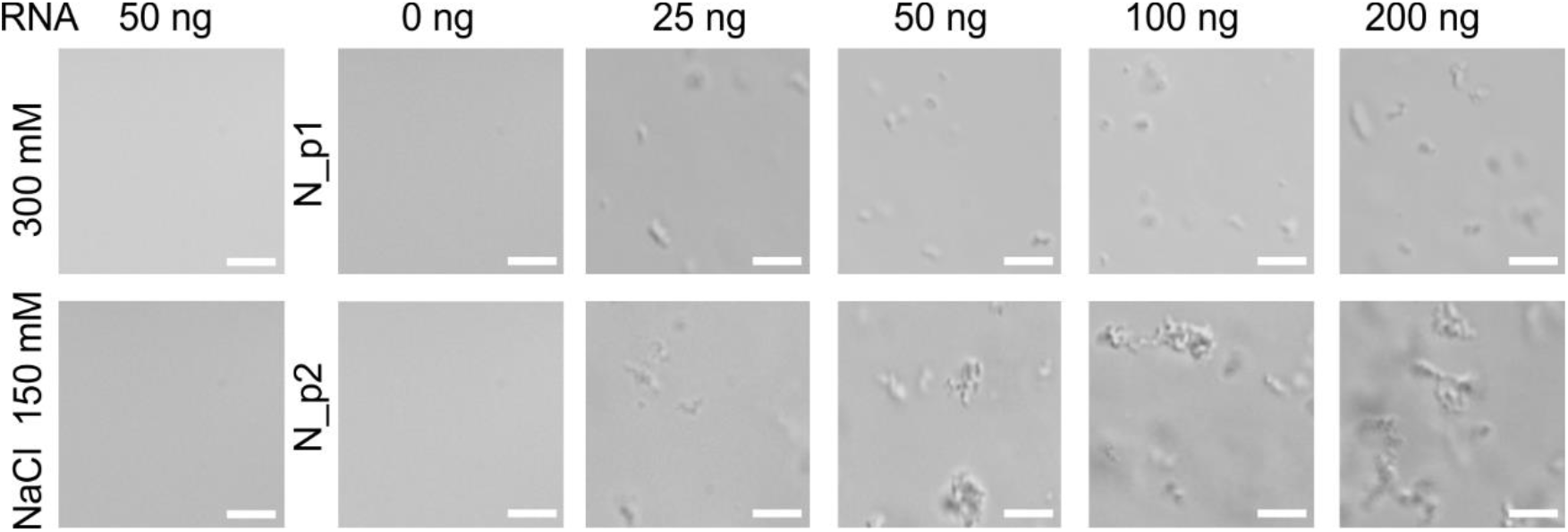
Formation of aggregates in the presence of bacterial RNA. Representative DIC images of the solutions containing 50 ng of bacterial RNA in a buffer containing 300 mM NaCl or 150 mM NaCl (first column from the left), or 2 µg of the N protein (1 mg/ml) in the presence of different amounts of RNA. The amount of used RNA is indicated at the top. The N_p1 was analysed in buffer containing 300 mM NaCl, and the N_p2 in buffer containing 150 NaCl. Scale bar - 5 µm.

A notable effect was observed on the drying tested samples. Several minutes after each sample preparation and microscopic examination, the buffer began to evaporate. This process was accompanied by the formation of relatively large round condensates. The N protein-containing condensates were persistent even when a sample evaporated and salt crystals were deposited (Figure 12A, B). These phenomena were observed in both the solutions that had and had not undergone LLPS (see Figure 10), as well as in the case of the N_p1 and N_p2 protein solutions supplemented with the bacterial RNA. As a control, we used FKBP39, which has previously been shown to induce LLPS. Once the phase-separated solution containing FKBP39 had dried, no condensates could be observed (Figure 12C). This observation suggested that the N protein-containing condensates were resistant to drying. We hypothesized that the obtained condensates are solid, which would suggest that the N protein drives not only liquid-liquid but also liquid-solid phase transitions. To test this hypothesis, we performed the FRAP experiment. The obtained results indicated that in the region of interest, after illumination with highly intense light, the fluorescence did not recover (Figure 12D). The fluorescence intensity measured over a period of 70 s was close to zero (Figure 12E). This observation strongly supported the hypothesis that SARS-CoV-2 can undergo other liquid-liquid phase transitions.

**Figure 12.**
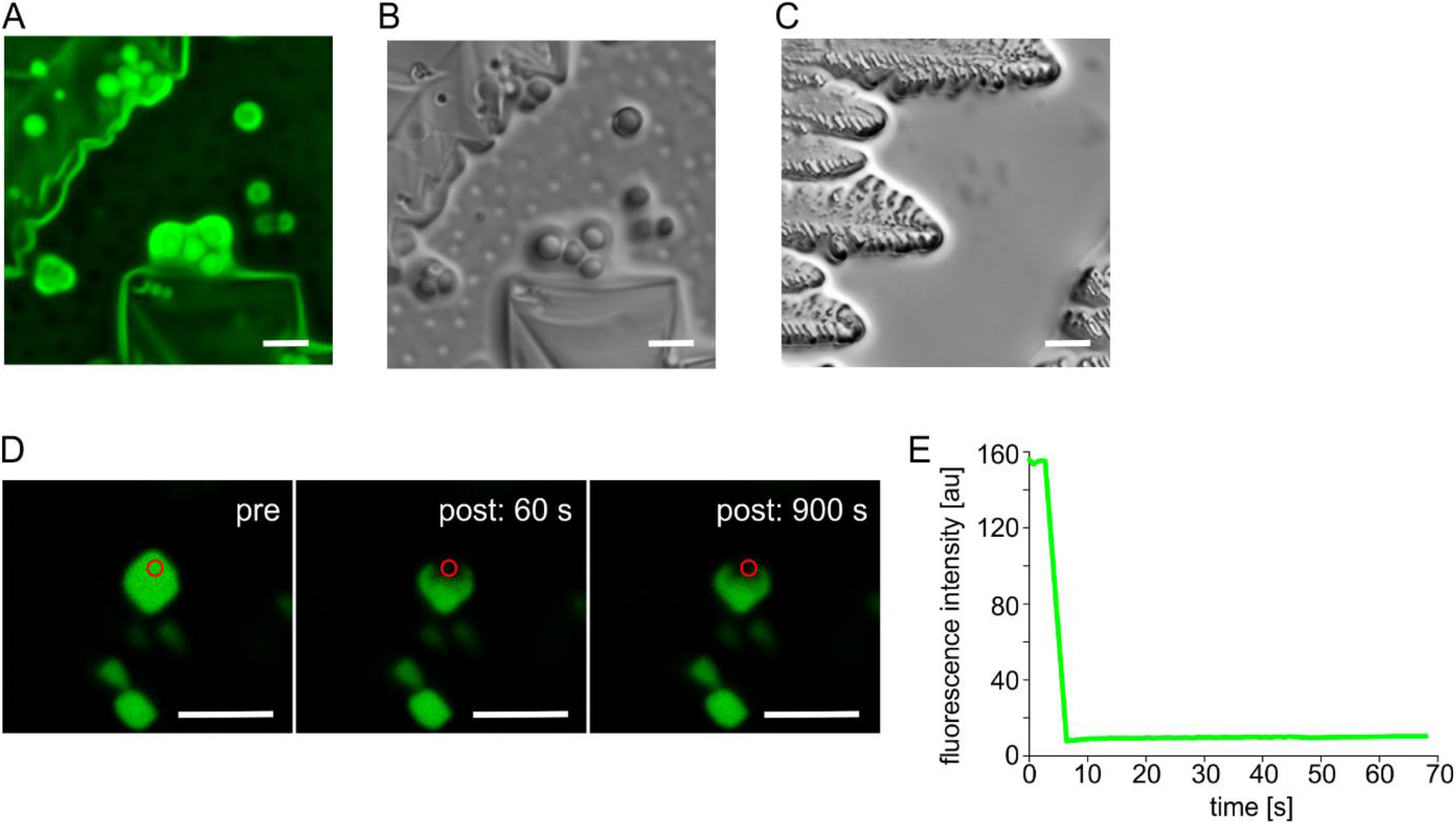
Condensates formed in a drying solution. The representative images of the condensed forms of the N protein labelled with ATTO 488 fluorescent dye. A) Condensates observed using widefield fluorescence microscopy; B) condensates observed using DIC microscopy; C) dried solution containingFKBP39 protein used as a control; D) confocal images obtained before (pre), 60 s and 900 s after the photobleaching (bleach time 2500 ms); E) FRAP curve of the fluorescently-labelled N protein over a 70 s period of time. Bleached area is indicated by red circle; a.u. – arbitrary units, Scale bar - 5 µm.

## Discussion

Coronavirus SARS-CoV-2 biology has been intensively studied since the start of the global pandemic, which began suddenly at the end of 2019. This pandemic has affected and disturbed nearly all aspects of human life on a worldwide scale. Strategies to eliminate the virus by developing an effective therapy or protective vaccine are in demand. The N protein, the most abundant protein in the virion (Kang *et al*, 2020; Grifoni *et al*, 2020), appears to be a promising target for such treatments. Furthermore, this protein is a determinant of the virulence and pathogenesis of SARS-CoV-2 (Yasui *et al*, 2008), which is considered a highly immunogenic antigen (Burbelo *et al*, 2020). The primary structure of the N protein is highly conserved and not vulnerable to mutations, which is especially interesting in the context of the appearance of new SARS-CoV-2 variants that are resistant to some of the newly developed “spike protein-targeted” vaccines (Liu *et al*, 2021)(Emary *et al*, 2021). Recently, reports focusing on the N protein as a potential vaccine target for SARS-CoV-2 were published (Oliveira *et al*, 2020; Grifoni *et al*, 2020; Bhattacharya *et al*, 2020; Rakib *et al*, 2020; Chukwudozie *et al*, 2020; He *et al*, 2021). For this reason, the characterizing the molecular basis of the functionality of this protein is highly important.

To date, a number of papers characterizing the structure, function and molecular properties of the N protein have been published. However, based on our results, N protein samples obtained according to previously described purification methods (Zeng *et al*, 2020; Perdikari *et al*, 2020; Ye *et al*, 2020; Zinzula *et al*, 2020; Lu *et al*, 2020) might have been severely contaminated with nucleic acids from the recombinant expression host organisms. In fact, during the preparation of this manuscript, the problem of randomly bound nucleic acids from *E. coli* serving as the host organism to the N protein was reported (Tugaeva *et al*, 2020; Zeng *et al*, 2020). According to the authors, the complete elimination of nucleic acid impurities was challenging (Tugaeva *et al*, 2020). In this study, we attempted to develop an effective purification method that enables the elimination of all nucleic acids bound to the N protein. We first tested the most commonly used procedures that are known to disrupt the interactions of protein-nucleic acids, e.g., treatment with a buffer with high ionic strength. However, the interactions between the protein and nucleic acids are very strong (Zeng *et al*, 2020), and this approach was determined to be insufficient. Even treatment with 2.5 M sodium chloride did not result in the dissociation of the nucleic acids from the protein. Further analyses of various chromatographic techniques and experimental conditions allowed the determination of the optimal combination of methods (see Figure 2) that enable the full-length N protein expressed in a prokaryotic system to be effectively and efficiently purified. Importantly, since nucleocapsid proteins share similar features and present a strong inherent ability to bind nucleic acids, the problem of their contamination by nucleic acids can be seen as not limited to the protein from the SARS-CoV-2 virus but should also be considered in the case of nucleocapsid proteins from other viruses. As we are convinced that DNA/RNA contamination may affect the properties of the protein, updated biochemical characterization of the molecular properties of the SARS-CoV-2 N protein is urgently important.

The performed comparative biochemical analysis of the N protein purified to homogeneity (protocol developed in this study) and the protein in complex with nucleic acids from *E. coli* revealed notable differences in structural properties. Although the shapes of the CD spectra curves indicated that the protein remained partially disordered in both samples, differences in the quaternary structure were observed. The N protein enables viral RNA packaging (Masters, 2019; McBride *et al*, 2014) and is thus a key player during the self-assembly of new viral particles. Therefore, studies aiming to determine the assembly of the N protein into higher-order RNP complexes are important. In previous reports, the formation of dimers and higher oligomers by the SARS-CoV-2 N protein was reported (Zeng *et al*, 2020; Perdikari *et al*, 2020). Notably, the CTD domain of the protein has been suggested to form an octamer that is maintained by electrostatic interactions (Perdikari *et al*, 2020). Our results do not support the formation of high oligomers by the full-length protein. The N protein purified according to the protocol developed in this study formed only homodimers, with higher oligomers not being observed. Notably, the findings presented by Chang et al. for the SARS-CoV N protein are in keeping with our results (Chang *et al*, 2006). The control experiments performed for the N protein sample purified according to the standard protocol, i.e., containing randomly bound nucleic acids, revealed the existence of dimeric species, as well as higher-molecular-mass complexes. Importantly, it is still not known whether high-molecular-mass complexes are higher oligomers of the N protein (Ye *et al*, 2020) or whether this protein’s dimers carry nucleic acids. Solving this puzzle may help to elucidate the mechanism underlying the viral genomic material’s self-assembly and viral particle packaging.

LLPS has recently been intensively studied in the context of the functionality of the SARS-CoV-2 N protein. It appears that this thermodynamically driven process initiates the formation of condensates in a host cell, which are the precursor of virion particles. Further research attempting to elucidate this mechanism may enable the biology of the virus to be better understood and the target points in the viral replication cycle to be identified. This research could be useful in the context of developing new treatment strategies. In general, liquid condensates were shown to be formed via transient interactions (Brangwynne *et al*, 2015)(Hyman *et al*, 2014). Previous studies revealed that condensates formed by the SARS-CoV-2 N protein contain viral RNA, as well as other components, including host proteins (Tugaeva *et al*, 2020; Lu *et al*, 2020; Savastano *et al*, 2020; Perdikari *et al*, 2020). The recruitment of host proteins is an important factor in altering host cell metabolism. Establishing a method to enable the selective dissolution of SARS-CoV-2 liquid condensates or the prevention of their formation may facilitate efforts to identify a means of combating viral infection.

In this study, we made important observations concerning the propensity for the spontaneous formation of dense condensates by the N protein. Previously, the N protein was reported to form liquid condensates in the presence of different types of RNA (Lu *et al*, 2020). We did not observe such an effect under the conditions used in our study. However, we did observe the formation of liquid condensates formed by the N sample contaminated with nucleic acids at low ionic strength. It was shown that the N protein can bind a fragment of viral genomic RNA encoding the N protein (Iserman *et al*, 2020a)(Carlson *et al*, 2020). Therefore, it is possible that the N protein expressed in bacterial cells may be complexed with specific mRNA encoding N, rather than random prokaryotic sequences. For both the N_p1 and N_p2 samples, we were able to observe the appearance of some filamentous precipitates after the addition of the total bacterial RNA. What is important is that phosphorylation of the N protein was recently proposed to cause liquid-like droplets, while nonphosphorylated N proteins, which correspond to the unmodified N protein expressed in bacteria, were also shown to form amorphous filamentous aggregates in the presence of RNA (Carlson *et al*, 2020).

A notable effect, though not fully understood, was observed when the tested solutions containing the N protein were dried. This phenomenon was observed for both N protein forms, i.e., the one that was free of nucleic acids and the one that was in a complex with bacterial nucleic acids. Upon drying, we observed the formation of condensates, which did not fuse together and were even persistent when the water completely evaporated and salt deposits became visible. The morphology, behaviour and minimal molecular diffusion kinetics suggested that these condensates were not liquid condensates but other more ordered forms, e.g., gel-like or solid. Notably, SARS-CoV and SARS-CoV-2 are the only known coronaviruses with N proteins that do not possess cysteine residues, which are believed to be responsible for enhancing the stabilization of other coronavirus virions by forming disulfide bonds (Surjit *et al*, 2004b). Therefore, we hypothesize that the more organized structures observed in this analysis may facilitate the compact character of RNP and stabilize the SARS-CoV and SARS-CoV-2 protein structures.

In our review of the literature, we found the study of Stewart et al. (Stewart *et al*, 2017), which concerns the sandcastle worm adhesive system. Images documenting the phase separation of secreted glue surprisingly resemble images presenting the phenomenon described above, which was observed during the drying of the samples. Notably, the phase transition of the sandcastle glue from a fluid adhesive into a porous solid adhesive joint was shown to be induced by substantial pH differences between the secretory system (pH below 6) and seawater (pH over 8), as well as changes in its ionic composition (Stewart *et al*, 2017). Importantly, it was documented that changes in pH associated with cellular uptake via endocytosis can be used to trigger the disassembly of the coacervate-based delivery vehicle (Blocher & Perry, 2017). In fact, the entry of avian coronavirus infectious bronchitis virus (IBV) into cells was stopped by inhibitors of pH-dependent endocytosis (Chu *et al*, 2006). These findings motivated us to speculate about the putative adhesive properties of the N protein and its potential biological significance. Naskalska et al. (Naskalska *et al*, 2019) showed that the S protein of the human coronavirus NL63 was not necessary for virus adhesion to host cells. Therefore, an alternative method of cell penetration independent of the S interaction with ACE2 was proposed by Naskalska et al. (Naskalska *et al*, 2019). These researchers suggested viral entry into the host cell by M protein interactions with heparan sulfate proteoglycans (HSPGs) (Naskalska *et al*, 2019). Notably, the studies were performed in the presence of the N protein. Considering that the N protein tightly interacts with M (He *et al*, 2004; Lu *et al*, 2021), we can assume that additional experiments should be performed to elucidate the role of the N protein in the proposed alternative cell penetration pathway. Notably, Blocher and Perry (Blocher & Perry, 2017) discussed chirality in addition to pH, ionic strength and temperature as a factor influencing phase separation. Importantly, in this study, we showed that the N protein possesses a unique optically active chiral centre, which might be involved in such regulation.

At present, the phenomenon of the formation of solid condensates, as discussed in this report, is not fully understood and warrants further investigation. Nevertheless, the results of this study indicate that the phase separation driven by biomacromolecules has some unexplained aspects and that further research is necessary to understand its mechanisms and physiological significance.

## Materials and methods

### Protein expression and purification

The cDNA of the N protein, which was generously received from Prof. N.J. Krogan (Department of Cellular and Molecular Pharmacology (UCSF), Gladstone Institutes, San Francisco) was used as a template during PCR. The primers introduced restriction sites for NcoI and NotI endonucleases in forward and reverse primers, respectively (underlined in primer sequences): the forward primer sequence was 5’-cgcg<u>ccatgg</u>CGATGAGCGATAACGGCCCCC, and the reverse primer sequence was 5’-gcgc<u>gcggccgc</u>**<u>TTA</u>**CGCCTGAGTAGAATC GGC. The uppercase letters in the primer sequence represent the sequence that is present in the N protein. The stop codon is bolded. The purified PCR product was cloned into the pET-SUMO vector digested with NcoI and NotI restriction enzymes. The final construct (pET-SUMO/N) was confirmed by DNA sequencing.

*Escherichia coli* BL21(DE3) cells were transformed with the obtained pET-SUMO/N construct, plated on LB agar supplemented with 50 µg/ml kanamycin, and incubated at 37°C overnight. A single colony was used to inoculate 20 ml of LB medium containing 50 μg/ml kanamycin. The starting culture was incubated overnight at 37°C in a rotary shaker operated at 200 rpm. 15 ml of the starting culture was then used to inoculate 1 l ZYM 5052 of autoinduction medium (Studier, 2005) supplemented with 100 μg/ml kanamycin. The incubation was conducted at 37°C until the optical density at 600 nm (OD_600_) was 2.0. Finally, the protein was expressed at a reduced overnight temperature of 20°C. The culture was collected by centrifugation (4000×g, 20 min, 4°C), resuspended in 10 ml of lysis buffer (50 mM Tris-HCl, pH 8.0, 300 mM NaCl, 5 mM β-mercaptoethanol, 20 mM imidazole; supplemented with 0.2 mg/ml phenylmethylsulfonyl fluoride (PMSF), DNase I, RNase A and lysozyme) and lysed by sonication. The lysate was clarified by centrifugation (20 000 rpm, 50 min). The 6xHis-SUMO-N protein was present in the supernatant (soluble fraction). The supernatant pH was adjusted to 8.0 and incubated with Ni^2+^-NTA His-binding resin (Novagen) that was pre-equilibrated with binding buffer (50 mM Tris-HCl, pH 8.0, 300 mM NaCl, 5 mM β-mercaptoethanol, 20 mM imidazole). The resin was then loaded on a reusable column (20 ml, Clontech) and washed with 20 ml of binding buffer. After washing, the column was closed, and 10 ml of binding buffer, together with 0.5 mg SUMO hydrolase (dtUD1) provided by our department, was added to the resin. All the components were gently mixed and incubated overnight at 4°C. Finally, the digested N protein was collected as the flow through (FT). The SUMO protein and dtUD1 protease, as both remained on the resin, were separated from the N protein.

For the first purification method (protocol 1), N-containing fractions were concentrated on the Amicon Ultra-4 Centrifugal Filter Unit (Merck/Millipore; molecular weight cut-off 30.0 kDa) and loaded on a Superdex 200 10/300 GL size exclusion column (GE Healthcare Life Sciences) pre-equilibrated with 20 mM HEPES, pH 8.0, 500 mM NaCl and connected to the ÄKTAavant (GE Healthcare Life Sciences) system operated at 0.5 ml/min at room temperature. Selected fractions containing N protein (N_p1) were used for further experiments.

For the second purification method (protocol 2), N-containing fractions were concentrated on the Amicon Ultra-4 Centrifugal Filter Unit (Merck/Millipore; molecular weight cut-off 30.0 kDa) and loaded on two connected in series 1 ml Heparin-Sepharose columns (Cytiva) pre-equilibrated with 20 mM HEPES, pH 8.0, 500 mM NaCl, connected to the ÄKTAavant (GE Healthcare Life Sciences) system. Bound proteins were eluted with a NaCl concentration linear gradient (0.5-0.7 M) for 15 min. Selected fractions containing the N protein were concentrated and loaded on two connected Superdex 200 10/300 GL size exclusion columns (GE Healthcare Life Sciences) pre-equilibrated with 20 mM HEPES, pH 8.0, and 500 mM NaCl. Selected fractions containing the N protein (N_p2) were used for further experiments. All stages of chromatography were monitored by ultraviolet (UV) absorbance at 220, 260 and 280 nm wavelengths. The parameters of the N protein, either fused or not fused to 6×His-SUMO, were calculated using the ProtParam server (Gasteiger *et al*, 2005).

### Analytical size-exclusion chromatography (SEC)

Analytical SEC was conducted at room temperature on a Superdex200 10/300 GL column (GE Healthcare Life Sciences) connected to the ÄKTAavant (GE Healthcare Life Sciences) system operated at 0.5 ml/min. The UV absorbance at 260 and 280 nm was monitored. Protein purified according to protocol 1 or protocol 2 (1 mg/ml) was injected in a volume of 0.2 ml.

### SEC-MALS

50 μl of the N protein samples (1 mg/ml) purified according to protocols 1 or 2 were loaded onto a Superdex 200 Increase 10/300 column (GE Healthcare) pre-equilibrated with 20 mM HEPES, pH 8.0, 500 mM NaCl buffer, connected to the OMNISEC - GPC/SEC (Malvern) system, and operated at 0.5 ml/min. Elution was monitored by in-line detectors: refractive index (RI), light scattering (Right Angle Light Scattering, RALS/Left Angle Light Scattering, LALS and Multi Angle Light Scattering, MALS) and a UV-PDA spectrometer. Data analysis and MM calculations were performed using OMNISEC software (Malvern).

### Detection of nucleic acids

To examine whether the N protein purified according to protocol 1 was contaminated with nucleic acids, the N_p1 samples were treated with anionic detergent at an elevated temperature. 10 µl of the N_p1 sample was incubated for 15 min at 50°C or supplemented with 1 µl of 10% SDS solution and incubated for 15 min at 50°C. To verify the ability of increased ionic strength to disrupt the integration between the N protein and prokaryotic nucleic acid, the N_p1 samples were mixed with buffer supplemented with various concentrations of sodium chloride. The samples were incubated for 30 min at room temperature. To identify the type of nucleic acids bound to the N protein, the protein samples at a concentration of 1 mg/ml were incubated with 50 µg of RNase A or DNase I (Sigma Aldrich). The reaction was carried out for 30 min at room temperature.

### Agarose gel electrophoresis

The prepared samples were analysed electrophoretically on agarose gels. The samples were mixed with 6× loading buffer and loaded onto a 1% agarose gel prepared using 1× TBE buffer. As a nucleic acid marker, 1 kb or 100 bp DNA ladders (Thermo Scientific) were used. The gel contained 0.005% Simply Safe (EURx) nucleic acid dye. The electrophoresis was run at a constant voltage of 90 V. The gel was visualized using the GelDoc Imaging System (Bio-Rad).

### Qualitative analysis on the N protein

For the qualitative analysis of the samples obtained at each stage of the purification procedure, electrophoretic and spectroscopic analyses were conducted. Electrophoresis was performed under denaturing conditions on 12% polyacrylamide gels (1 mm thickness) developed in a Tris/glycine system (Laemmli, 1970) at a constant amperage of 20 mA/gel. As a protein weight marker, an UPMM (Thermo Fisher Scientific Inc., USA) was used. After protein separation, the gel was stained with BlueStain Sensitive (EURx) dye following the manufacturer’s instructions (https://eurx.com.pl/docs/specs/en/e0298.pdf). After destaining, acrylamide gel was visualized using the GelDoc Imaging System (Bio-Rad). For the spectroscopic examinations, 2 µl of the studied protein sample was loaded onto a zeroed NanoDrop spectrophotometer (Thermo Scientific). The absorption spectra were recorded in the range of 220-350 nm.

### SV-AUC

Sedimentation velocity analytical ultracentrifugation (SV-AUC) experiments were carried out using a ProteomeLab XL-I analytical ultracentrifuge (Beckman Coulter) equipped with an AN-60 Ti 4-hole rotor, 12 mm path-length charcoal-filled double sector epons, and quartz windows. Scans were recorded using absorbance optics. The experiments were performed at a temperature of 20°C and at a rotor speed of 40 000 rpm. The protein was analysed at three concentrations of 0.25 mg/ml, 0.5 mg/ml and 0.75 mg/ml in 20 mM HEPES buffer that contained 500 mM NaCl (pH 7.2). The volume of each sample was 400 µl. A series of 150 scans was obtained at 1.5 min intervals. The collected scans were fitted to a continuous size-distribution *c(s)* model implemented in SEDFIT version 14.1 software (Brown & Schuck, 2006)(Schuck, 2000). Buffer density (*ρ* = 1.0121 g/cm^3^) and viscosity (*η* = 0.010447 mPa×s) were calculated from the composition of the buffer using SEDNTERP software (Harding, 1992). A partial, specific volume of 0.71919 was calculated from the amino acid sequence using SEDNTERP (Schuck, 2000).

### Circular dichroism (CD)

CD spectroscopy experiments were conducted using a JASCO J-715 spectropolarimeter. The final spectra were obtained as an average of three measurements. Each measurement was conducted at a scanning speed of 20 nm/min, a data resolution of 1.0 nm, and a band width of 1.0 nm. The experiments were performed at 20°C using a Peltier-type temperature controller. The measurements were performed in a 1 mm path-length quartz cuvette using N_p1 and N_p2 samples at a concentration of 10 µM in 20 mM HEPES buffer containing 500 mM NaCl and with a pH of 7.2. The spectra were collected in a spectral range of 200–340 nm. The obtained spectra were corrected for the contribution of the buffer, and the measurements were converted to molar residual ellipticity units (Kelly *et al*, 2005). Deconvolution of the averaged CD spectrum was performed using CDNN 2.1 software (http://www.gerald-boehm.de/) using molar ellipticity values from 210-260 nm.

### Phase separation assay

For microscopic examinations, the N_p1 and N_p2 samples in 20 mM HEPES, 500 NaCl, pH 7.2 buffer were concentrated to 10 mg/ml using the Amicon Ultra-4 Centrifugal Filter Unit (Merck/Millipore; molecular weight cut-off 30.0 kDa). The concentrated solution was used to prepare a series of working solutions. The N_p1 and N_p2 samples were analysed without additives at various concentrations ranging from 1-10 mg/ml in buffer containing 500 mM NaCl. Next, the N_p1 sample at a concentration of 1 mg/ml was analysed in 20 mM HEPES buffer containing different concentrations of NaCl ranging from 100–500 mM. The N_p2 sample was analysed analogically but at concentrations of 1 mg/ml and 2 mg/ml. Both N protein preparations were analysed in the presence of total prokaryotic RNA. For fluorescence microscopy, the N protein was labelled with ATTO 488 dye (Sigma Aldrich) according to the manufacturer’s instructions. Widefield microscopy was performed using an Axio Observer 7 (Carl Zeiss) inverted microscope with 100 × oil and 1.3 numeric aperture objective lenses. Differential interference contrast (DIC) and fluorescence images were collected with an Axiocam 305 colour camera (Carl Zeiss).

### FRAP experiments

For FRAP experiments, 3D images of fluorescently labelled protein condensates were acquired using a Leica TCS SP5 II confocal system equipped with argon and helium-neon lasers, a 63x oil and 1.4 numeric aperture objective lens. ATTO 488 dye was excited using 488 nm light, and emitted fluorescence was observed in the range of 500-535 nm. A series of FRAP experiments were performed using a parked light beam from an argon laser (210 µW). Bleaching times varied from 100 ms to 2500 ms.

### *E. coli* BL21 RNA purification

Total bacterial RNA was purified using a total RNA mini kit (A&A Biotechnology) according to the manufacturer’s protocol.

## Acknowledgements

We thank Prof. N.J. Krogan (Department of Cellular and Molecular Pharmacology (UCSF), Gladstone Institutes, San Francisco) for kind gift of the N protein cDNA.

This work was supported by a subsidy from The Polish Ministry of Science and High Education for the Faculty of Chemistry of Wroclaw University of Science and Technology. Confocal fluorescence microscopy imaging was supported by the Polish National Science Center (NCN) 2017/27/B/NZ3/01065.

## Author contributions

Conceptualization: BGM; Experiments and data analysis: AT and MKA; Initial cDNA cloning experiments: BGM; FRAP experiments: MZ; Methodology: AT, MKA, BGM, AO, MZ; Resources: BGM, AO, JD; Visualization: AT, MKA, BGM; Manuscript writing: AT, MKA, BGM; Manuscript review and editing: AT, MKA, BGM, AO, MZ, JD;

## Expanded View Figure legends

**Figure EV1.**
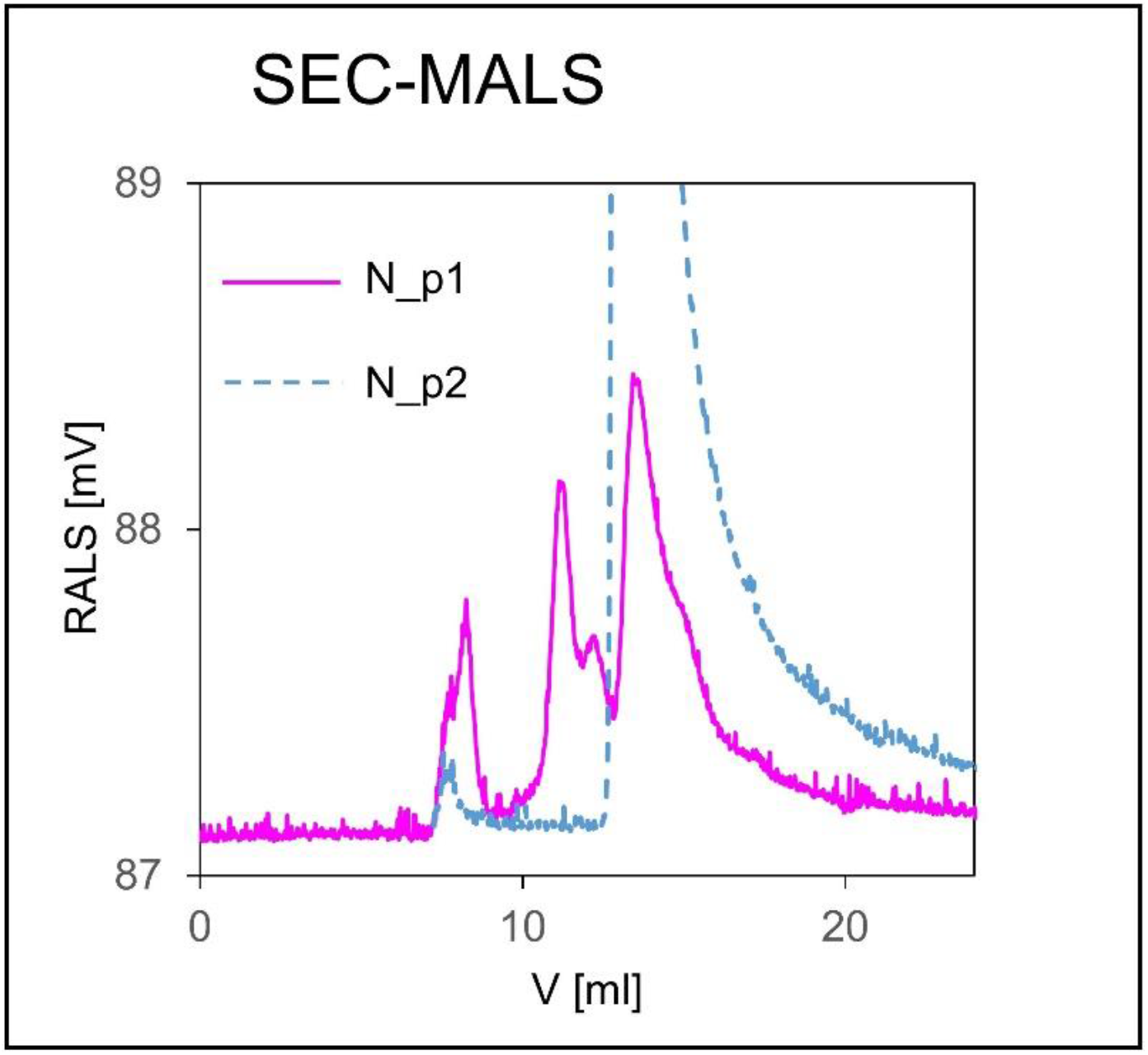
The SEC-MALS profile of the N_p1 sample. The SEC-MALS profile of the N_p1 sample (magenta line) was superimposed on the SEC-MALS profile of the N_p2 sample (dashed blue line).

